## Supplemental Figs S1-S12 and Table S1 for "*daf-16*/FOXO blocks adult cell fate in *Caenorhabditis elegans* dauer larvae via *lin-41*/TRIM71"

### Supplemental Materials

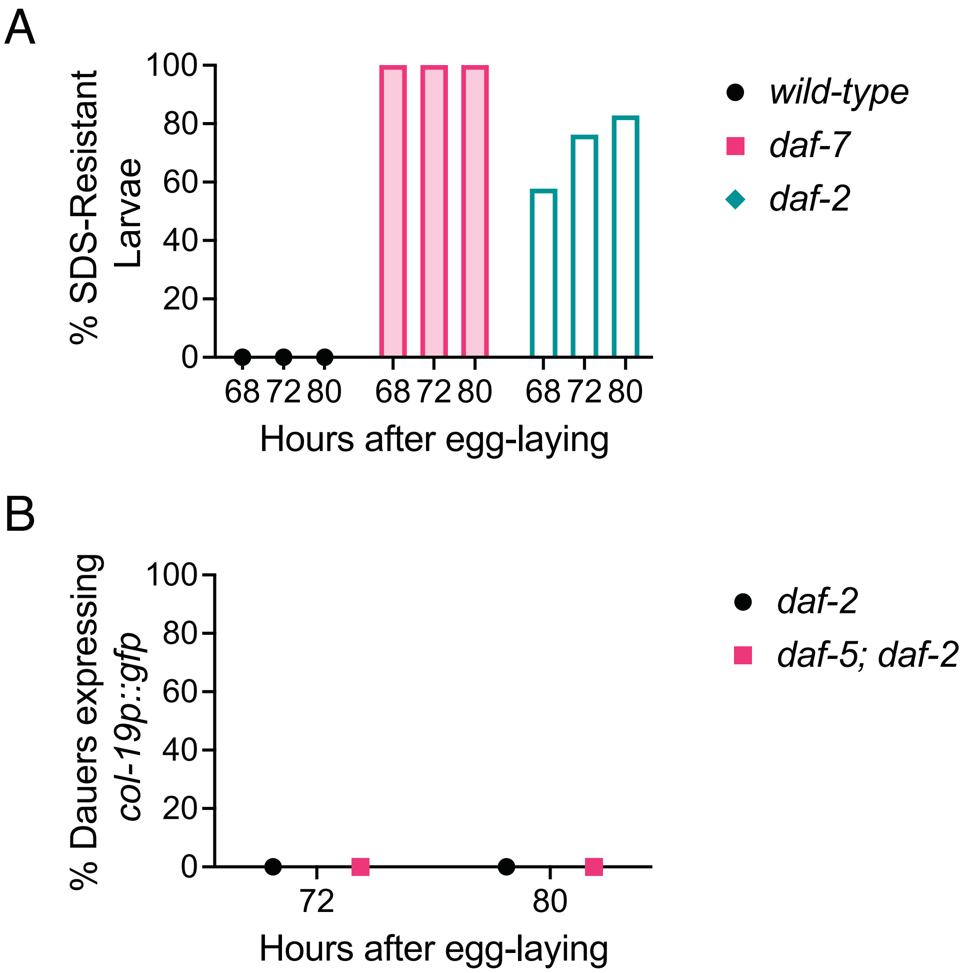


**Figure S1. *daf-5* is not required to block *col-19::gfp* expression during dauer.** The *daf*-7/TGFβ pathway regulates dauer formation in parallel to the insulin-like pathway (Fielenbach and Antebi 2008; Baugh and Hu 2020). *daf-5* encodes a Sno/Ski protein that works in a complex with DAF-3/SMAD to regulate transcription downstream of DAF-7/TGFβ signaling (Graca *et al.* 2004). *daf-5(0)* mutants are dauer-defective, but can be induced to enter dauer when combined with the *daf-2(e1370)* allele (Vowels and Thomas 1992; Larsen *et al.* 1995). (A) *daf-2(e1370)* mutants develop slowly and acquire dauer characteristics approximately one day later than wild-type or *daf-7(e1372)* larvae (Ruaud *et al.* 2011; Nika *et al.* 2016). Since *daf-16; daf-7* dauer larvae express *col-19p::gfp* most penetrantly within one day of dauer entry, we used SDS-resistance to determine a time that might correspond to "early dauer" in this strain. SDS-resistance is acquired at the end of the L2d-to-dauer molt and continues throughout dauer (Cassada and Russell 1975). n = 67-91. (B) Percent of dauer larvae expressing *col-19p::gfp.* This expression was never observed in either control or *daf-5(0)* dauer larvae at times when many early dauer larvae should have been present. n = 26-54.

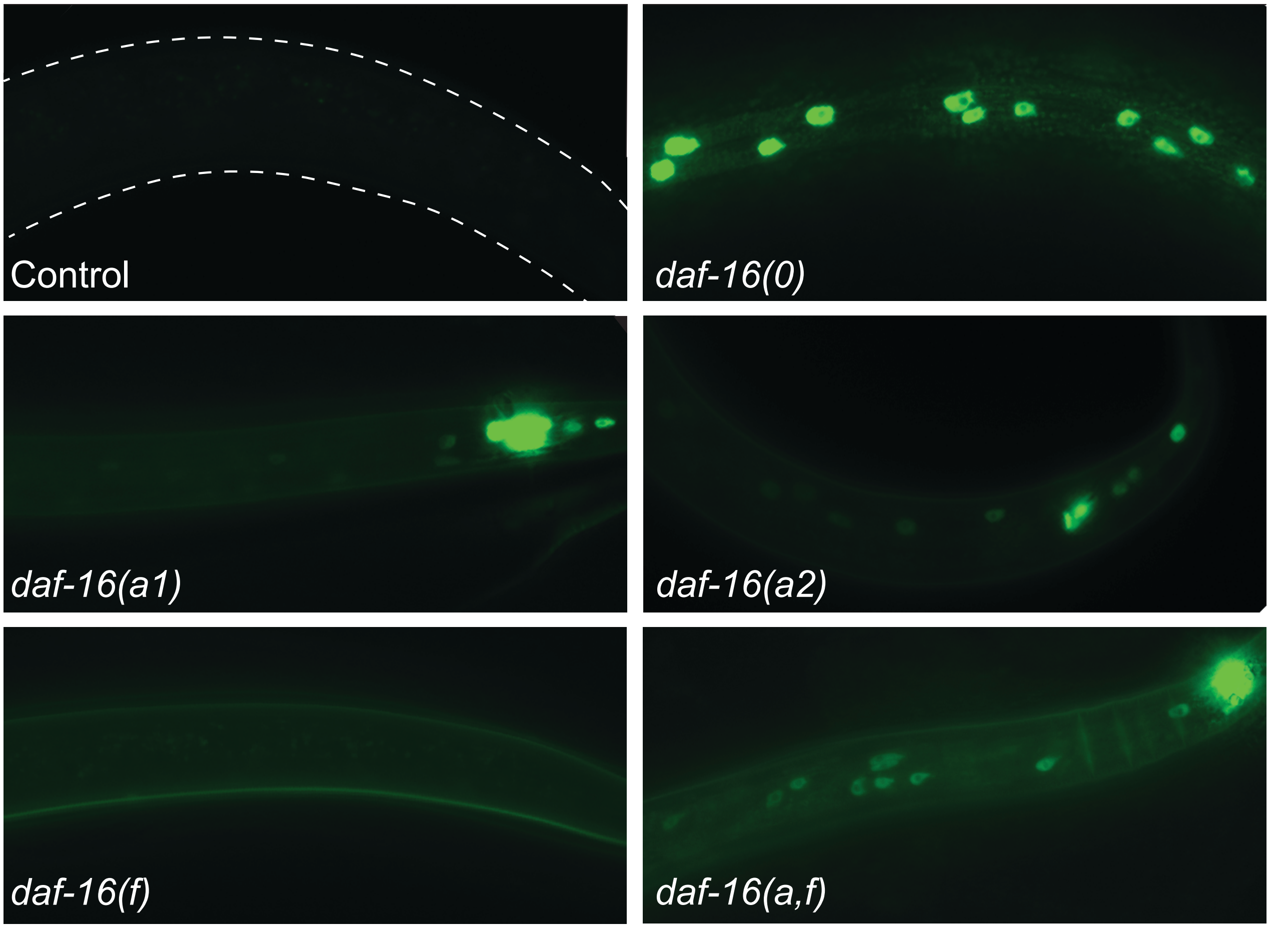


**Figure S2. Dauer larvae lacking 1-2 isoforms of *daf-16* display *col-19p::gfp* more dimly and in fewer cells than *daf-16(0).*** Representative micrographs of dauer larvae scored in Fig 1D. In Fig 1D, any *col-19p::gfp* expression in the lateral hypodermis was considered “on”, however there are qualitative differences in that expression in the different strains. Dashed lines in the control indicate the boundaries of the dauer larva, as determined by DIC optics.

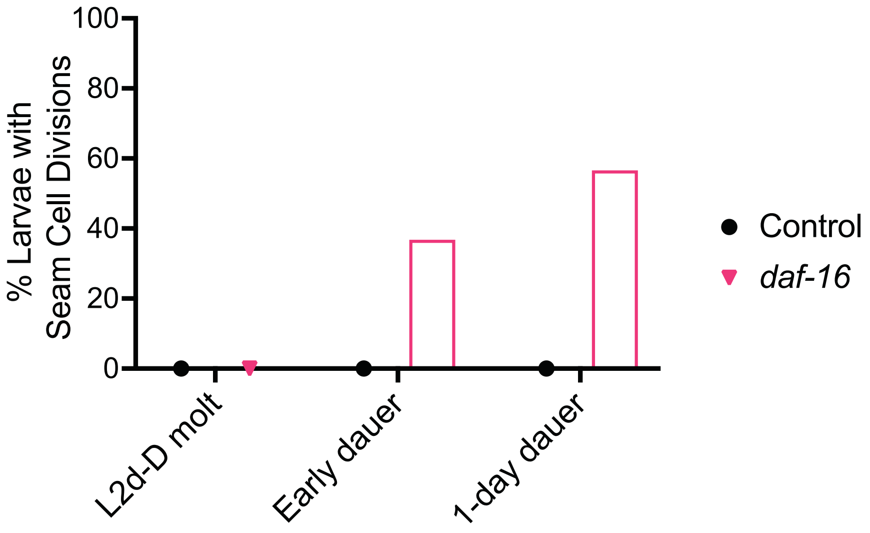


**Figure S3. Seam cells divide and remain unfused in *daf-16(0)* dauer larvae.** Percent of dauer larvae with one or more seam cells actively dividing at the time of scoring. Seam cell divisions were identified based on *ajm-1::gfp* expression from the *wIs78* transgene (Abrahante *et al.* 2003). *ajm-1::gfp* is a translational fusion that labels apical junctions. *wIs78* also contains *scm::gfp* that labels seam cell nuclei (Koh and Rothman 2001). Junctions were present between all seam cells in both control and *daf-16(0)* dauer larvae, however cell divisions were only seen in *daf-16(0)* dauer larvae. n = 10-87.

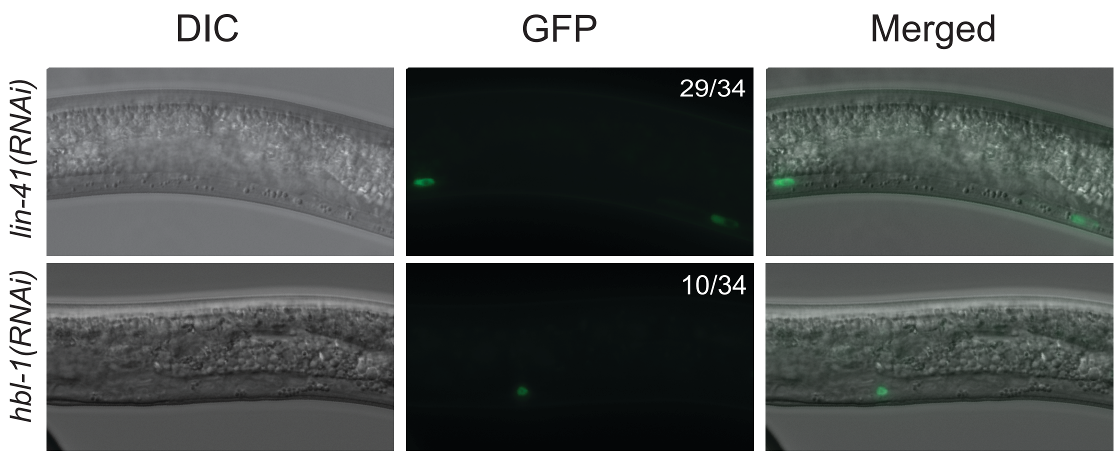


**Figure S4. RNAi of *hbl-1* and *lin-41* induces *col-19p::gfp* expression in non-hypodermal tissues.** Numbers indicate larvae expressing *col-19p::gfp* in at least one cell in the relevant tissue over the total number of dauer larvae scored. (Top) *lin-41* RNAi induces *col-19p::gfp* expression in vulva precursor cells (VPCs) during dauer at high penetrance. The larva shown expressed *col-19p::gfp* in P5.p and P7.p, but overall which VPCs expressed *col-19p::gfp* varied from worm to worm and did not correlate with the predicted cell fate they will adopt. The exposure time in the fluorescence channel was 200ms. (Bottom) *hbl-1* RNAi induces *col-19p::gfp* expression in an unidentified non-hypodermal cell during dauer at lower penetrance. The exposure time in the fluorescence channel was 50ms.

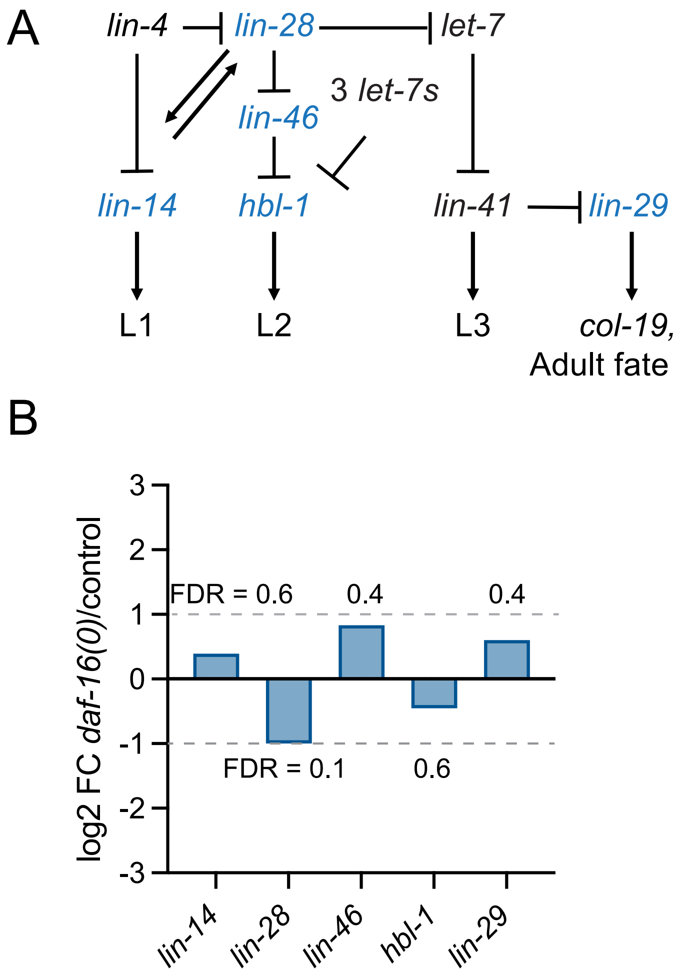


**Figure S5. Levels of most core heterochronic genes are essentially unaffected by *daf-16* during dauer.** (A) The network of heterochronic genes that regulates stage-specific seam cell fate during continuous development. “3 *let-7s”* indicates the *let-7* sisters, *mir-48, mir-84,* and *mir-241.* All protein-coding genes except *lin-41* are indicated in blue. (B) $Log_2$ fold change of the protein-coding heterochronic genes other than *lin-41,* comparing *daf-16(0)* to control dauer larvae. $Log_2$ fold change was calculated by DESeq2 from mRNA-seq data. Dashed lines indicate 2x fold upregulation or downregulation. False discovery rates (FDR) are indicated for each gene.

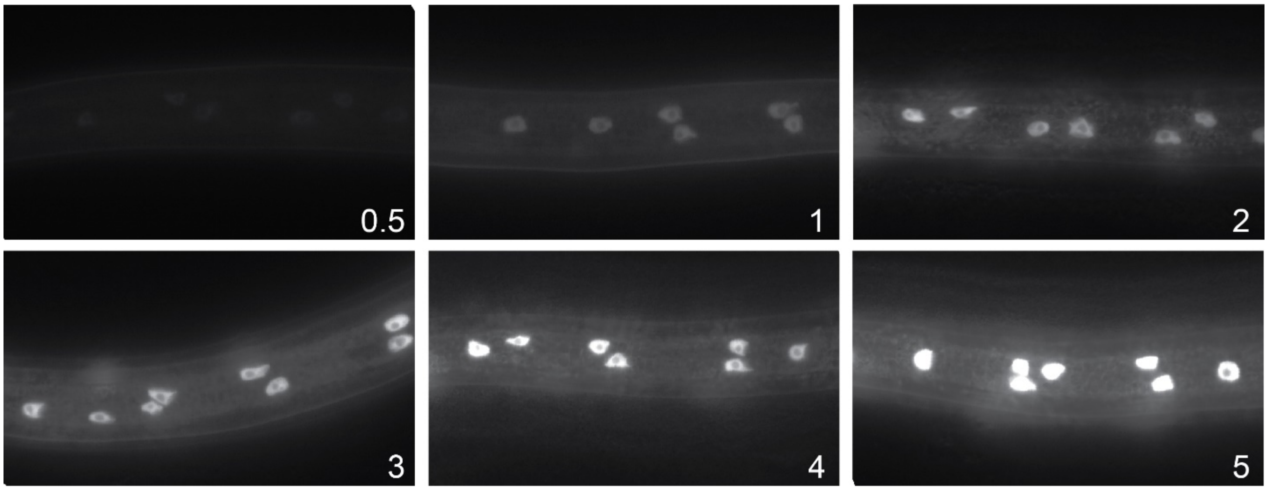


**Figure S6. Fluorescence scale used to determine relative fluorescence intensity of *col-19p::gfp*.** Fluorescence images of *col-19p::gfp* in dauer larvae were taken from a strain we found to produce a wide range of expression levels, XV160, *daf-7(e1372); maIs105; unk-1(xk6).* These images were taken using compound microscopy and identical settings, including an exposure time of 125ms. The images were then exported as 8-bit tiffs for ImageJ analysis. We identified six representative images distributed across the range of ImageJ values (0-250). We assigned numbers (0.5-5) to these images that reflect their distribution. A value of 0.25 was used if expression was visible, but less than that in the 0.5 panel. The fluorescence scale values with ImageJ values in parenthesizes were as follows: 0.5 (22), 1 (45), 2 (104), 3 (149), 4 (200), and 5 (250). These six images were then used as a scorecard to compare to experimental images for figures 3B, 4A, and 5B. For these experiments, a 63x objective was used to take 2-3 images along the length of the dauer larva, using identical settings for all strains being compared. Each image was then subjectively compared to the fluorescence scale and assigned a value 0-5, with 0.5 values being used if an image was between two whole-number images. The values for each worm were averaged together to create an overall score for the worm.

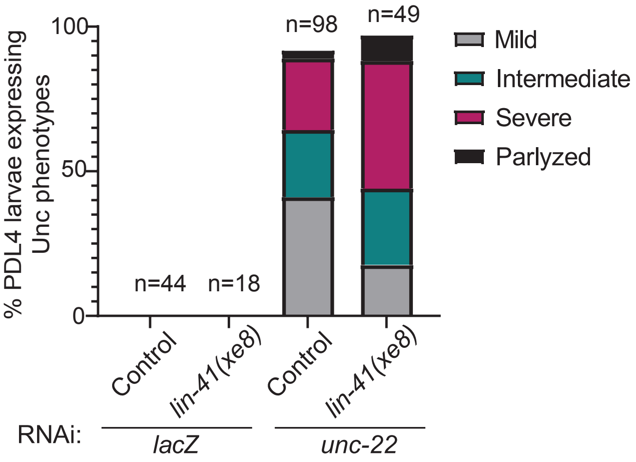


**Figure S7. The *lin-41(xe8)* strain is not defective in RNAi**. Unc phenotypes produced by *unc-22(RNAi)* were assessed during forward locomotion and binned into categories Mild: occasional twitching that did not interrupt movement. Intermediate: occasional twitching that did interrupt movement. Severe: constant twitching but still capable of forward locomotion. Paralyzed: worms whose twitching was so severe that they were not able to move forward. *unc-22* RNAi experiments were carried out in parallel to the experiments shown in figure 3. However, because dauer larvae did not display strong Unc phenotypes, dauer larvae were moved to new RNAi plates at 20˚C and allowed to recover to the post-dauer L4 stage. Numbers indicate the total number of post-dauer L4 larvae assessed.

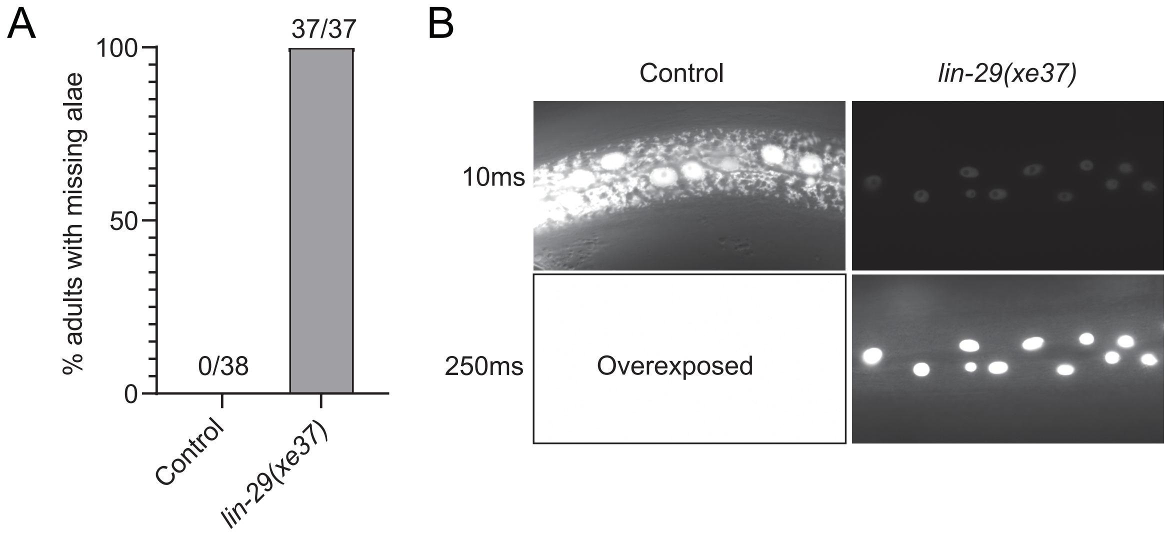


**Figure S8. The *lin-29(xe37)* strain displays reiterative phenotypes in adults.** (A-B) The background for all strains was *daf-7(e1372); maIs105[col-19p::gfp]*. (A) *lin-29(xe37)* mutant adults do not have adult alae. Numbers indicate the number of worms with alae defects over the total number of adults. (B) *lin-29(xe37)* mutant adults express very low levels of *col-19p::gfp*. Control worms exposed for 250ms (bottom, left) were too overexposed to determine the boundaries of the worm. Representative images are shown (n = 37-38).

A

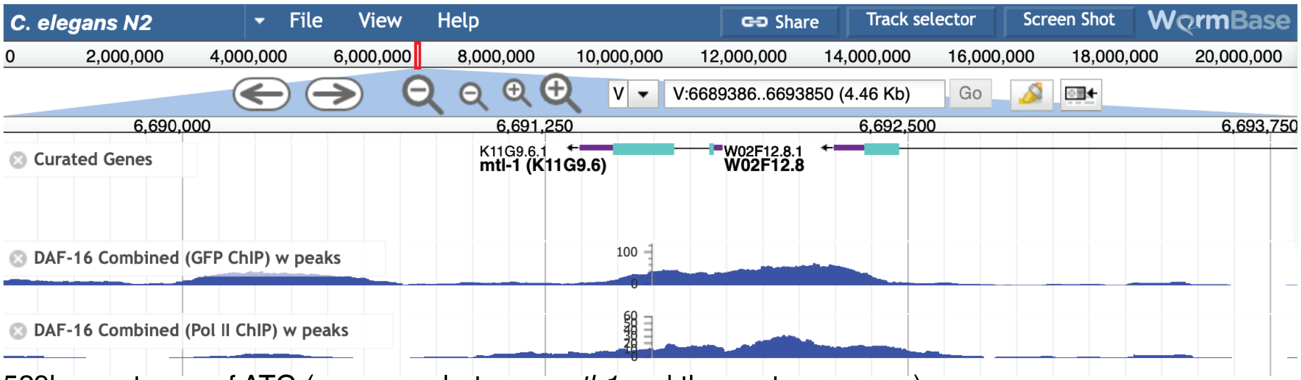


529bp upstream of ATG (sequence between *mtl-1* and the upstream gene):

atggcttgcccaatgagcaagtgaacttttcccacgtctactactaaattattgtcgatctataattctttcgcttatc
aatcttgtcaattgaaataaaacaatatttttctaattctttttggaacgaacacacgtgtttaaatgaatgttgtgct
aaaaacgtcacatcaatggtacgtgaatgttgcaaacaccttgtcaataactgataaaatcagaaactagagctgtgac
tgaatcgtatactagaacggagtctctctaaaaacgttctaaaaacaaacaaaaaattgtgcaaaggattagagtgctc
aagatcaatgagcaaactcacaatcaactatctgttattgttttgggtctcttctatatctctattcttttagtatcat
aagtttgtacattgtgacagggccaccctcttttatcacatatttgaagtg**gtaaaca**gccagcaaaaaccaaataaaa
aggcagtgagaaaaagaagaaggcagctcaatttgactgctgaaattaagaaatcATG

B

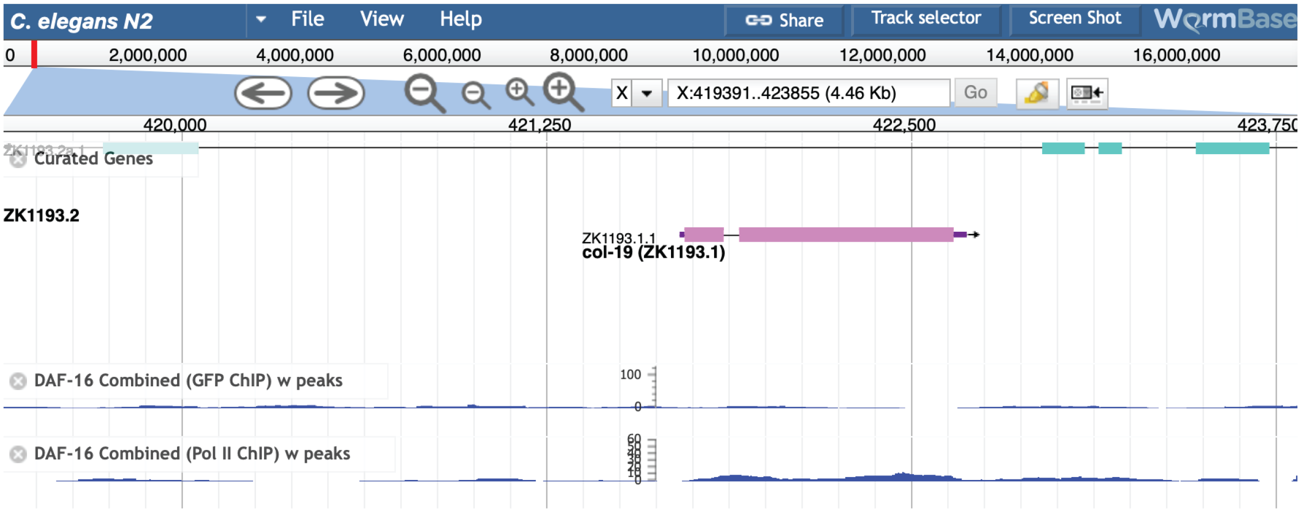


846bp upstream of ATG (promoter sequence included in *col-19p::gfp*):

aagcttccaaacgtccctattaggaaatgtaaacttattcccaaaaaattaaaaattccagagaaagtagacaaatttc
agaaaacttaccgcgctacatctaatactttctcataggttttttattgggaaactgatgaaaattatttgaattcat
aataaaataaattcgtatttagcatttgaaaatttgcaccaatgtattatttaattttttttttcgaaaatttaacgca
ttttctctctaaaaactcgaaatttagtgtgttctaaacaacagtaagcatacaatacctttgttcaaaattgacgtgc
tttctgaaccaatatggttagtttcaaaaattttttgtattataggatagaaatatttggaaataaattttaaaaccaaa
cttatgcctttctcttttagtatcccagctaggtaatattttagtatttgcccaaatccttgaagtaaggagtatataa
tttttgaaaaacaataaaactccagataattcatagtttttttctcgaaagaaaatttttgagattagttattgaacttc
atttttgaacattattcgttgaaaaacactcgctttgtctttattttcaaaaaaattccgatttccccaaccagaaaaa
aaaacagatagaagaaatttctccttaaatttcattgtccatctctcttggaaacacattatctatcaaatgaaaaacg
catttttttttcctggcagaaaaatgaaattggttagattacactggttaggtttgaaggtgtaactttcgctttctcag
caactttcagtataaaaggaaacggtcaccatttagaaagacatcagttcatcaacATG

**Figure S9. DAF-16 is unlikely to bind the *col-19* promoter to regulate transcription.** (A, top) Screenshot of Genome Browser from WormBase (WS280) showing 4.46kb of chromosome V, centered on *mtl-1,* a confirmed transcriptional target of DAF-16 (Barsyte *et al.* 2001; Li *et al.* 2008). The tracks display modENCODE ChIP data for DAF-16 binding. Clear peaks are observed upstream of *mtl-1.* (A, bottom) DNA sequence beginning immediately 3' to W02F12.8, the gene upstream of *mtl-1.* This sequence likely contains promoter elements that regulate *mtl-1* expression. A canonical DAF-16-Binding Element (DBE, red) is located 76bp upstream of the *mtl-1* ATG. (B, top) Screenshot of Genome Browser from WormBase (WS280) showing 4.46kb of chromosome V, centered on *col-19.* No DAF-16 ChIP peaks are visible. (B, bottom) The 846bp upstream of the *col-19* ATG which comprises the regulatory sequence driving *col-19p::gfp* expression (Liu *et al.* 1995). No canonical DBEs (GTAAACA or TGTTTAC) (Furuyama *et al.* 2000) were found in this sequence.

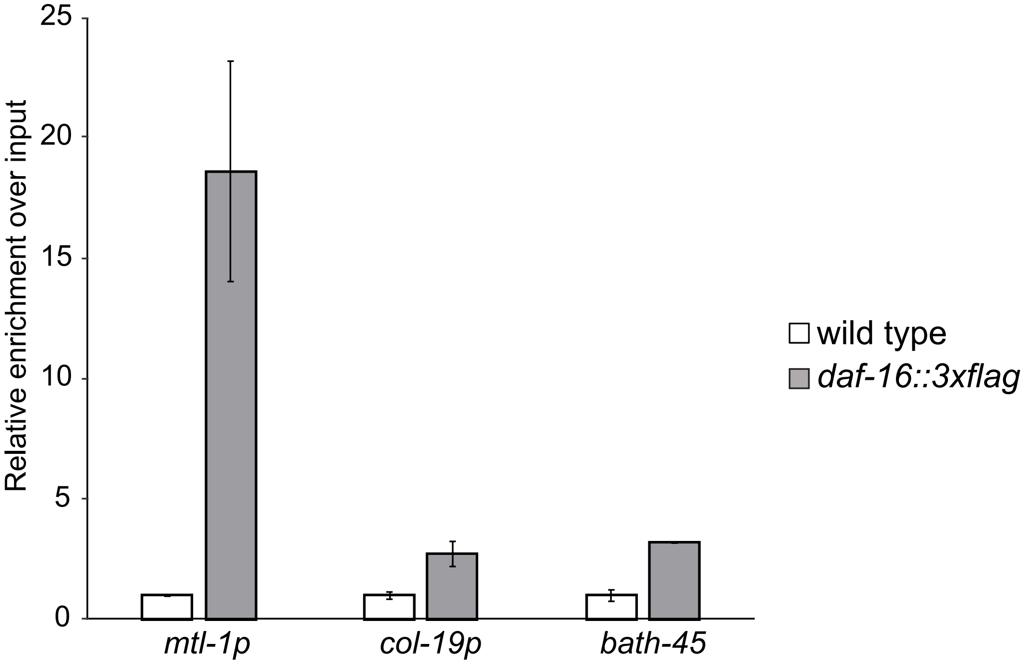


**Figure S10. DAF-16 is not enriched at the *col-19* promoter**. ChIP-qPCR experiments were performed on N2 or *daf-16(ar620[daf-16::zf1-wrmScarlet-3xFLAG])* dauer larvae. Binding of DAF-16-3xFLAG was first normalized to input, and then to the average of the respective wild-type value (mean +/- SD for two technical replicates) is shown. Binding to the *col-19* promoter was not observed, whereas there was substantial binding to the promoter of the known DAF-16 target *mtl-1* (Barsyte *et al.* 2001; Li *et al.* 2008). The coding region of the heterochromatinized gene *bath-45* was used as negative control. This figure shows the second of two biological replicates; the first replicate is shown in figure 6A.

A

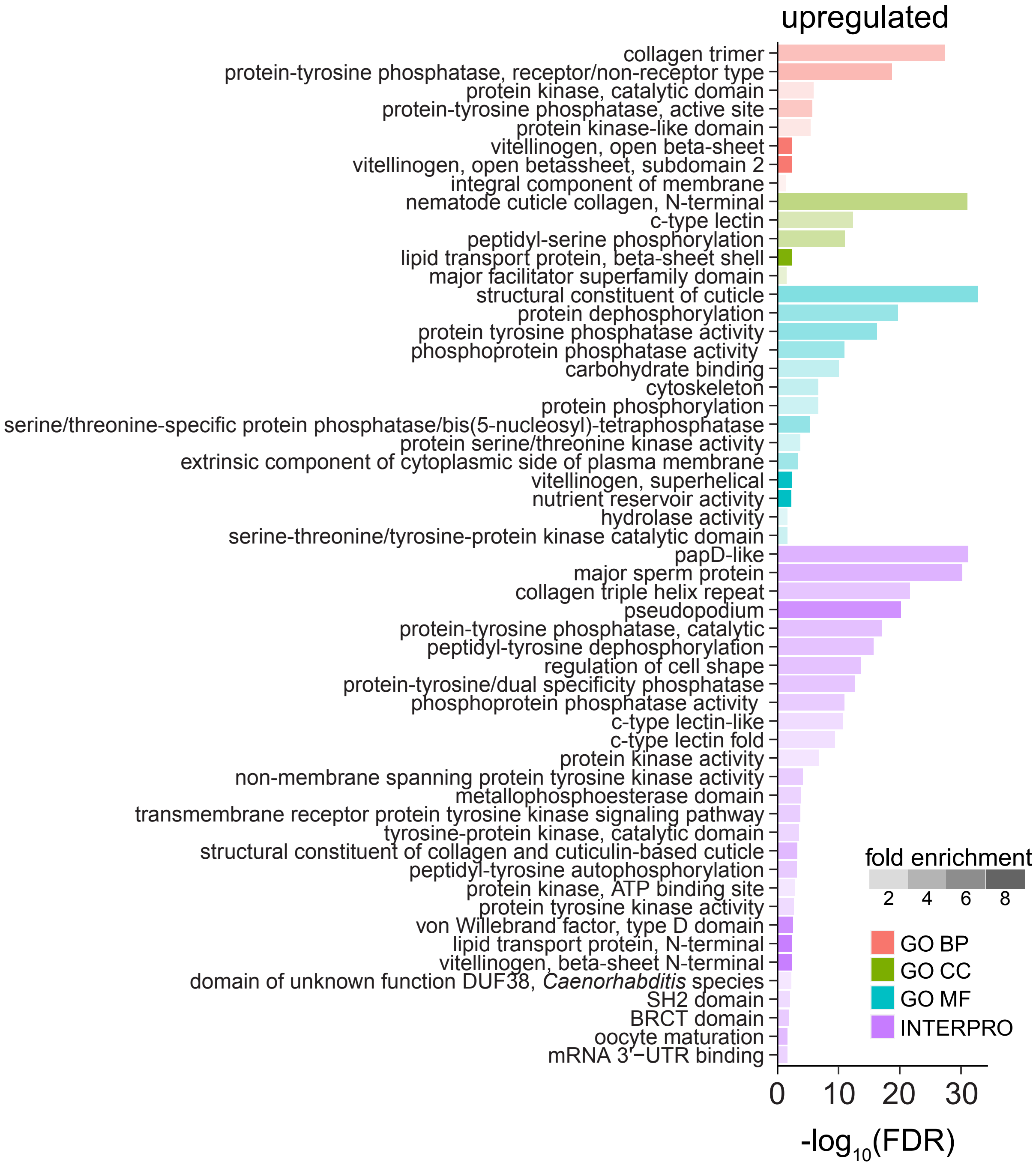
upregulated
collagen trimer
protein-tyrosine phosphatase, receptor/non-receptor type
protein kinase, catalytic domain
protein-tyrosine phosphatase, active site
protein kinase-like domain
vitellinogen, open beta-sheet
vitellinogen, open betassheet, subdomain 2
integral component of membrane
nematode cuticle collagen, N-terminal
c-type lectin
peptidyl-serine phosphorylation
lipid transport protein, beta-sheet shell
major facilitator superfamily domain
structural constituent of cuticle
protein dephosphorylation
protein tyrosine phosphatase activity
phosphoprotein phosphatase activity
carbohydrate binding
cytoskeleton
protein phosphorylation
serine/threonine-specific protein phosphatase/bis(5-nucleosyl)-tetraphosphatase
protein serine/threonine kinase activity
extrinsic component of cytoplasmic side of plasma membrane
vitellinogen, superhelical
nutrient reservoir activity
hydrolase activity
serine-threonine/tyrosine-protein kinase catalytic domain
papD-like
major sperm protein
collagen triple helix repeat
pseudopodium
protein-tyrosine phosphatase, catalytic
peptidyl-tyrosine dephosphorylation
regulation of cell shape
protein-tyrosine/dual specificity phosphatase
phosphoprotein phosphatase activity
c-type lectin-like
c-type lectin fold
protein kinase activity
non-membrane spanning protein tyrosine kinase activity
metallophosphoesterase domain
transmembrane receptor protein tyrosine kinase signaling pathway
tyrosine-protein kinase, catalytic domain
structural constituent of collagen and cuticulin-based cuticle
peptidyl-tyrosine autophosphorylation
protein kinase, ATP binding site
protein tyrosine kinase activity
von Willebrand factor, type D domain
lipid transport protein, N-terminal
vitellinogen, beta-sheet N-terminal
domain of unknown function DUF38, *Caenorhabditis* species
SH2 domain
BRCT domain
oocyte maturation
mRNA 3'−UTR binding
0
10
20
30
-log10(FDR)
fold enrichment
2
4
6
8
GO BP
GO CC
GO MF
INTERPRO

B

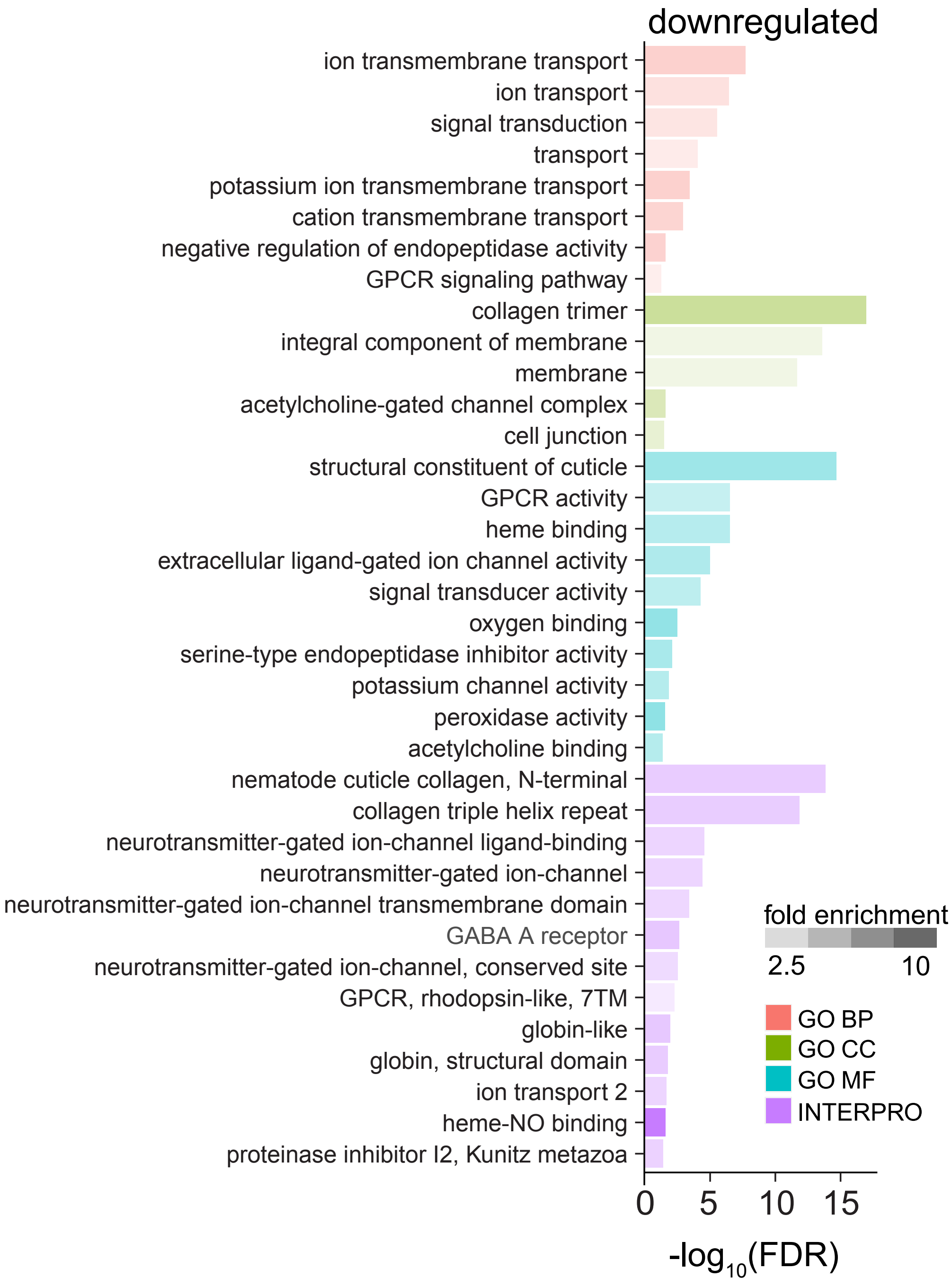


**Figure S11. Functional annotation clustering.** DAVID analysis showing gene ontology (GO) and InterPro terms that were significantly enriched (p ≤ 0.05, Bonferroni corrected) in genes whose expression changed ≥2x (FDR ≤ 0.05) in *daf-16(0); daf-7* dauer larvae vs. *daf-7* (control) dauer larvae. Genes that were upregulated in *daf-16(0)* dauers are shown in (A); genes that were downregulated are shown in (B). GO BP = GO term biological process; GO CC = GO term cellular compartment; GO MF = GO term molecular function, INTERPRO = InterPro protein classification.

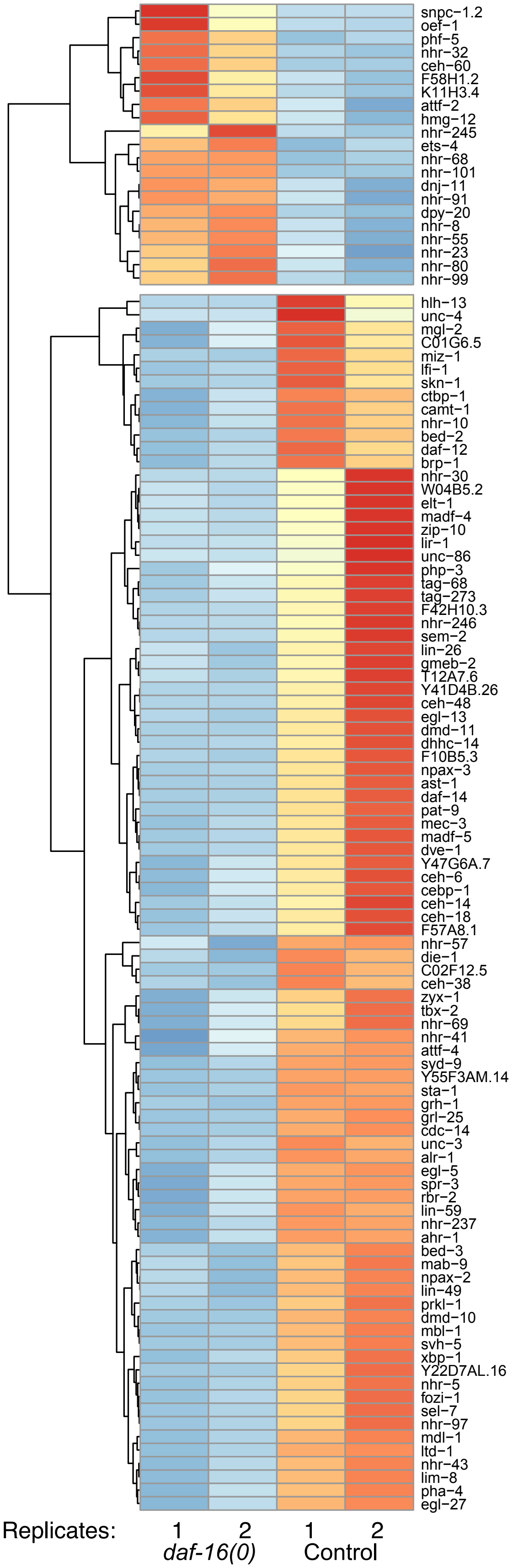


**Figure S12. Transcription factors differentially expressed in *daf-16(0)* dauer larvae compared to control dauer larvae.** Genes encoding transcription factors were defined based on their WormBase annotation (WS279). Although annotated as transcription factors, we noticed that the WormBase list also contained genes encoding RNA-binding proteins. We therefore subtracted genes identified as encoding RNA-binding proteins from the transcription factor list. From among these genes, we searched our mRNA-seq data to find those whose expression changed ≥2x (FDR ≤ 0.05) by DESeq2 analysis. This analysis produced 112 genes that fit our criteria. Those genes are shown here as heat maps depicting relative expression of these genes in *daf-16(0); daf-7* dauer larvae or *daf-7* (control) dauer larvae from mRNA-seq data. The row-normalized reads for each of two biological replicates per strain are depicted.

| Figure | Strain | Genotype | References |
|---|---|---|---|
| Fig 1A | VT1777 | *daf-7(e1372); maIs105[col-19p::gfp]* | *maIs105:* (Feinbaum and Ambros 1999) |
| | XV36 | *daf-16(mgDf50); daf-7(e1372); maIs105* | *mgDf50:* (Ogg *et al.* 1997) |
| | XV156 | *daf-16(mu86); daf-7(e1372); maIs105* | *mu86:* (Lin *et al.* 1997) |
| Fig 1B | VT1777 | *daf-7(e1372); maIs105* | |
| | XV36 | *daf-16(mgDf50); daf-7(e1372); maIs105* | |
| Fig1C | XV33 | *maIs105* | |
| | VT1750 | *daf-16(mgDf50); maIs105* | |
| Fig 1D | VT1777 | *daf-7(e1372); maIs105* | |
| | XV36 | *daf-16(mgDf50); daf-7(e1372); maIs105 (= daf-16(0))* | |
| | XV72 | *daf-16(tm3050); daf-7(e1372); maIs105 (= daf-16(a1))* | *tm3050, tm3052, tm6659:* (Chen *et al.* 2015) |
| | XV73 | *daf-16(tm3052); daf-7(e1372); maIs105 (= daf-16(a2))* | |
| | XV71 | *daf-16(tm6659); daf-7(e1372); maIs105 (= daf-16(f))* | |
| | XV74 | *daf-16(mg54); daf-7(e1372); maIs105 (= daf-16(a,f))* | *mg54:* (Ogg *et al.* 1997) |
| Fig 1E | CB1372 | *daf-7(e1372)* | |
| | VT2317 | *daf-16(mgDf50); daf-7(e1372)* | |
| Fig 2B | VT1777 | *daf-7(e1372); maIs105* | |
| Fig 3A | CB1372 | *daf-7(e1372)* | |
| | VT2317 | *daf-16(mgDf50); daf-7(e1372)* | |
| Fig 3B | VT1777 | *daf-7(e1372); maIs105[col-19p::gfp]* | |
| | XV181[1] | *lin-41(xe8)/lin-41(bch28[Peft3::gfp::h2b::tbb-2 3'UTR] xe70[∆3'UTR]); daf-7(e1372); maIs105* | *xe8:* (Ecsedi *et al.* 2015). *bch28 xe70:* (Aeschimann *et al.* 2019) |
| Fig 4A | VT1777 | *daf-7(e1372); maIs105[col-19p::gfp]* | |
| | XV253 | *lin-29(xe37); daf-7(e1372); maIs105* | *xe37:* (Aeschimann *et al.* 2019) |
| Fig 4B | VT1777 | *daf-7(e1372); maIs105* | |
| | XV238[2] | *lin-41(n2914)/nIs408[lin-29::mCherry, ttx-3p::gfp]; daf-7(e1372); maIs105* | *n2914:* (Slack *et al.* 2000) *nIs408:* (Harris and Horvitz 2011) |
| | XV245 | *nIs408; daf-7(e1372); maIs105* | |
| | XV239[2] | *lin-41(n2914)/nIs408; lin-29(n546); daf-7(e1372); maIs105* | |
| Fig 4C | HW1822 | *lin-29(xe61[lin-29::gfp::3xflag])* (for L3 experiments) | *xe61:* (Aeschimann *et al.* 2017) |
| | XV243 | *lin-29(xe61); daf-7(e1372)* (for dauer experiments) | |
| Fig 4D | XV243 | *lin-29(xe61); daf-7(e1372)* | |
| Fig 5A | XV36 | *daf-16(mgDf50); daf-7(e1372); maIs105* | |
| | XV87 | *daf-16(mgDf50); lin-29(n546); daf-7(e1372); maIs105* | |
| | XV254 | *daf-16(mgDf50); lin-29(xe37); daf-7(e1372); maIs105* | |
| Fig 5B | VT1777 | *daf-7(e1372); maIs105* | |
| | XV36 | *daf-16(mgDf50); daf-7(e1372); maIs105* | |
| | XV254 | *daf-16(mgDf50); lin-29(xe37); daf-7(e1372); maIs105* | |
| Fig 6A | N2 | wild-type | |
| | GS8924 | *daf-16(ar620[daf-16::zf1-wrmScarlet-3xFLAG])* | Gift from K. Luo, Greenwald lab |
| Fig 6B | CB1372 | *daf-7(e1372)* | |
| | VT2317 | *daf-16(mgDf50); daf-7(e1372)* | |
| Fig S1A | N2 | *wild type* | |
| | VT1777 | *daf-7(e1372); maIs105* | |
| | XV138 | *daf-2(e1370); maIs105* | |
| Fig S1B | XV138 | *daf-2(e1370); maIs105* | |
| | XV115 | *daf-5(m512); daf-2(e1370); maIs105* | *m512:* (Tewari *et al.* 2004) |
| Fig S2 | VT1777 | *daf-7(e1372); maIs105* | |
| | XV36 | *daf-16(mgDf50); daf-7(e1372); maIs105* | |

| Figure | Strain | Genotype | References |
|---|---|---|---|
| Fig S2 | XV72 | *daf-16(tm3050); daf-7(e1372); maIs105 (= daf-16(a1))* | |
| | XV73 | *daf-16(tm3052); daf-7(e1372); maIs105 (= daf-16(a2))* | |
| | XV71 | *daf-16(tm6659); daf-7(e1372); maIs105 (= daf-16(f))* | |
| | XV74 | *daf-16(mg54); daf-7(e1372); maIs105 (= daf-16(a,f))* | |
| Fig S3 | XV27 | *daf-7(e1372); wIs78[scm::gfp, ajm-1::gfp]* | *wIs78:* (Koh and Rothman 2001; Abrahante *et al.* 2003) |
| | XV29 | *daf-16(mgDf50); daf-7(e1372); wIs78[scm::gfp, ajm-1::gfp]* | |
| Fig S4 | VT1777 | *daf-7(e1372); maIs105* | |
| Fig S5 | CB1372 | *daf-7(e1372)* | |
| | VT2317 | *daf-16(mgDf50); daf-7(e1372)* | |
| Fig S6 | XV160 | *daf-7(e1372); maIs105; unk-1(xk6)* | |
| Fig S7 | VT1777 | *daf-7(e1372); maIs105* | |
| | XV81 | *lin-41(xe8)/lin-41(bch28 xe70); daf-7(e1372); maIs105* | |
| Fig S8 | VT1777 | *daf-7(e1372); maIs105* | |
| | XV253 | *lin-29(xe37); daf-7(e1372); maIs105* | |
| Fig S10 | N2 | wild-type | |
| | GS8924 | *daf-16(ar620[daf-16::zf1-wrmScarlet-3xFLAG])* | |

**Table 1. List of strains used in this study.**

1) Strain XV181 was maintained as heterozygotes. In this case, the relevant homozygous progeny were isolated the day before the experiment. Homozygous progeny were recognizable by lack of GFP expression from *bch28.*
2) Strains XV238 and XV239 were maintained as heterozygotes. For experiments individual gravid hermaphrodites were cloned out to several 60 mm plates and allowed to lay embryos for 3-5 hours at 24˚C. Parents were then removed. Plates where 100% of larvae expressed *ttx-3p::gfp* were discarded. From the remaining plates, all dauer larvae were scored for both *ttx-3p::gfp* expression (indicating *nIs408)* and *col-19p::gfp* expression.
